# Differential Expression Analysis for Metatranscriptomics with Sample-Paired Metagenomics Data

**DOI:** 10.1101/2022.12.08.519567

**Authors:** Binghao Yan, Di Wu

## Abstract

1

**Motivation:** Large microbial communities have contributed to essential ecosystems and multiple health and disease processes. Metatranscriptomics (MTX) is becoming increasingly important for profiling the gene expression pattern and functional activities of microbial communities. A fundamental task for analyzing the high-throughput sequencing data is the Differential Expression (DE) analysis, which aims to identify up/down-regulated genes/pathways across multiple conditions. However, DE analysis is complicated by the variation of underlying DNA copies, which is caused by the highly dynamic nature of microbial communities. The non-independent change between MTX and MGX may lead to false discoveries about which genes are biologically differentially expressed. Nevertheless, We can settle this problem by using the information from paired metagenomics (MGX) data.

**Results:** We proposed the MetaDePair, a statistical model for DE analysis in MTX with paired MGX data. It relied on a conditional Negative-Binomial distribution to model the count data. We showed that the adjustment of underlying DNA copies could significantly eliminate the variations of RNA data, thus improving the power of statistical inference. Also, we found that appropriate pre-filtering of zeros can also improve the sensitivity and precision of the model. Moreover, we proposed a method to simulate the paired MTX and MGX data. Using simulated data and real microbial data sets, we demonstrated that our method has a high statistical power while controlling the false discovery rate (FDR). We applied the tool to oral microbiome data and identified significant genes that are associated with Early Childhood Caries (ECC). Our tool enabled a more accurate and effective DE analysis based on MTX with paired MGX data and helped us improve the understanding of the functional characterization of microbial communities.

## 2 Introduction

Microbial communities have crucial roles in the ecological process and human health[1]. The high-throughput sequencing technique provides us with the ability to analyze a complex assemblage of microbes[2]. While metagenomics (MGX), which sequences the DNA extracted from the sample, can profile the taxonomic composition and genetic potential effectively, it does not directly reflect which gene or pathway is expressed or activated. Metatranscriptomics (MTX), which sequences the mRNA in the sample, enables the analysis of transcripts from genes that are truly expressed, thus elucidating gene expression patterns and the functional role of the microbial community. Previous studies about the gene expression of microbial communities have demonstrated the importance of using MTX to study gene expression dynamics under different conditions[3, 4, 5, 6]. At the same time, due to the uniqueness and complexity of metatranscriptome data, robust statistical pipelines are needed to provide reliable analysis results.

One fundamental question in analyzing MTX data is the differential expression (DE) analysis, which aims to identify microbial genes or pathways that are significantly differentially expressed in a microbial community. Various characteristics of microbial data have posed challenges to corresponding statistical frameworks. Firstly, the sequencing count or normalized data may be modeled as integers appropriately. The over-dispersion nature, which means that microbiome data is extremely noisy, reduces the power of statistical inference. Another factor that complicates the analysis of microbiome data is the excess of zeros[7], a problem that is critical in MTX and MGX data. The large proportion of zeros can be categorized into different origins: biological and technical. Biological zeros refer to genes that are lost or not expressed in the sample. In contrast, technical zeros represent zeros caused by various technical factors such as amplification bias or limited sequencing depths. Great attention should be paid to how to handle these zeros accurately.

However, given the success of the RNA-seq DE analysis method, there are few statistical workflows developed for MTX, and the analysis mainly relies on techniques borrowed from RNA-seq DE methods or microbiome differential abundance (DA) methods[8, 9, 10, 11, 12, 13]. Although single-organism RNA-seq and the microbial taxon abundance data share some common features with MTX data such as over-dispersion and sparsity, there are still gaps in data characteristics and model assumptions. One critical difference between MTX data and single-organism RNA-seq is the dependence of a gene’s RNA abundance on its underlying DNA abundance[14]. According to the central dogma, the variation in the abundance of encoding genes will possibly contribute to the changes in transcripts, and from a statistical perspective, introduce extra dispersions into the data. Thus, this underlying dependence between DNA and RNA will complicate the DE analysis. Only focusing on transcripts abundance may lead to false discoveries about non-regulatory changes since the observed changes may be caused by DNA abundance variation.

Methods have previously been proposed to tackle this underlying dependency between DNA and RNA abundance. For example, Klingenberg and Meinicke[15] propose a ‘taxon-specific scaling’ normalization method that explicitly accounts for variations in the taxonomic composition of transcripts. Specifically, their approach starts with assigning the feature profile to each taxon and then doing normalization within the taxon-specific count matrix. Finally, they recombine the normalized feature profile into a metatranscriptomics profile. Although this method eliminates changes caused by variations in taxonomic abundances, it has some drawbacks. Under some circumstances, the ‘taxon-specific scaling’ method will lose power since many genes in metatranscriptomics data can’t be assigned to a specific species or taxon. Also, it doesn’t consider other factors that will cause changes in gene copy numbers. When paired MGX-MTX data are available, directly considering the corresponding DNA abundance when analyzing MTX data can better eliminate the confounding. In another work[16], the author proposes a method to compute the “expression profile”, by calculating the log2-transformed ratio of transcript and gene abundance. This method decomposes the dissimilarity of MTX into two parts: the corresponding MGX and expression profiles. Biobakery 3[17] adds DNA as a covariate into the linear mixed model when identifying expression-level microbial metabolic biomarkers. A recent paper[14] evaluates six linear models for identifying DE, including models that directly take DNA into account. Although the methods above consider the baseline dependency between MTX and MGX, the simple linear model doesn’t address the special characteristics of microbiome data, thus facing the risk of losing power.

Here, we propose a model for differential expression analysis for metatranscriptomics with paired metagenomics data. It uses a conditional negative binomial distribution (see Methods) to describe the data. We demonstrate that our method, by using conditional estimation, can eliminate the extra variation introduced by DNA. We present a simple but effective way to filter the technical zeros in paired MGX-MTX data, which can increase the sensitivity of our model. Additionally, bootstrap is used for fold change shrinkage, which helps us decrease the false positives while not losing too much power. We furthermore compare our methods with existing DE tools using synthesized data and real datasets (Zero Out ECC-ZOE, and Inflammatory Bowel Diseases-IBD), showing that our methodology has a high statistical power while controlling the false discovery rate (FDR). Our method provides a new statistical workflow for identifying DE genes in MTX and offers the possibility for more accurate biological findings.

## 3 Methods

### 3.1 Description of Metatranscriptomics Datasets

We investigated the performance of our method using two metatranscriptomics datasets, both with paired metagenomics data. The first dataset (ZOE 2.0) was generated in a community-based epidemiological study of early childhood caries (ECC) in North Carolina, United States[18]. The second dataset was generated in a longitudinal study that investigated the association between gut microbial communities and IBD[19]. In the following two subsections, we described the studies and datasets in more detail and explained how we acquired the paired MGX-MTX data for subsequent analysis.

#### 3.1.1 The ZOE 2.0 study dataset

The ZOE 2.0 study mainly aimed to provide high-quality and large-scale information for research about pediatric oral health. The study collected clinical information on preschool-age children’s (ages 3-5) dental cavities and supragingival biofilm samples. A subset of participants’ biofilm samples (297/6404) had been carried forward with metagenomics and metatranscriptomics analysis, and the paired clinical information was available. After pre-processing, estimates of taxonomic composition, gene family, path abundance, and path coverage were produced from remained reads using HUMAnN2. There were a total of 204 distinct species and 402,937 distinct genes in the ZOE 2.0 metatranscriptome. We used all 297 samples with paired metagenomics and metatranscriptomics data for subsequent analysis. The ECC prevalence in these samples was 49% (147/297). We filtered genes with high zero proportions (>90%) and a total of 210158 genes were left.

#### 3.1.2 The IBD dataset

The objective of the IBD study was to understand the biological basis for inflammatory bowel diseases, which included Crohn’s disease and ulcerative colitis. The study followed 132 subjects for one year and generated profiles of multiple types of omics data, including metagenomics and metatranscriptomics. These gut microbiome data were in counts per million (CPM) and were derived using functional profiles 3.0 in HUMAnN 3.0. The datasets included a total of 1,595 metagenomics and 818 metatranscriptomics samples for 120 subjects.

A total of 109 subjects had both MGX-MTX data. Among them, one subject (Participant.ID: P6014) had no matched MGX-MTX data. We selected one paired MGX-MTX sample from each subject. Among these samples, one (Participant.ID: P6028) had extremely high zero proportions (99.68%) and we chose not to use it. We also needed to select genes that are both presented in MTX and MGX data. So we finally got a data matrix with 107 samples and 51519 genes for subsequent analysis.

### 3.2 Data preprocessing

There are cases that some genes are lost or not expressed. For example, some diseases may lead to the structural change of some proteins and thus inhibit the expression of certain genes. Here, we adopt a simple way to preprocess the data by filtering genes with zero proportions higher than 95%. The intuition behind this is that we tend to believe a gene with extremely high zero proportion is biologically not expressed.

### 3.3 Model and normalization

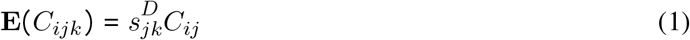

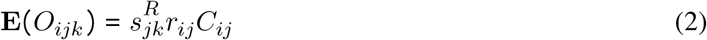

Since the expected abundance *C*_*ij*_ is not observed, we use a crust estimator 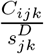 to replace it.

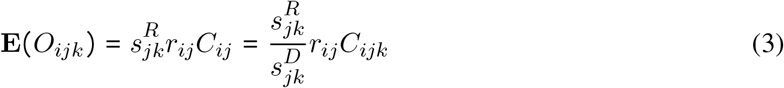

We then assume the read count *O*_*ijk*_ follows a negative binomial distribution conditional on *C*_*ijk*_,

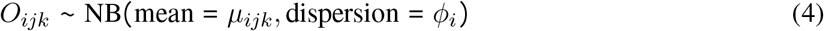

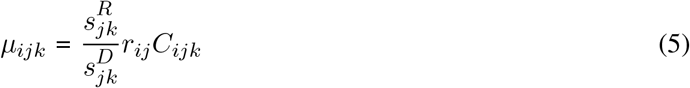

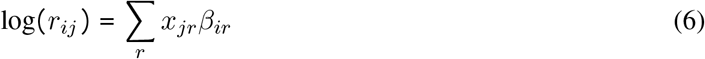

where *x*_*jr*_ is the group covariate and *β*_*jr*_ is the corresponding fold change. *ϕ*_*i*_ is the dispersion parameter and has the following relationship with the variance: var (*O*_*ijk*_) = *µ*_*ijk*_ (1 + *µ*_*ijk*_*ϕ*_*i*_). It’s important to note that the parameter we are interested in is the expression rate *r*_*ij*_, the absolute value of which represents the extent of regulatory change. We need to make statistical inferences on *β*_*ir*_ to identify differentially expressed genes.

There are various ways to estimate the normalization constants *s*_*jk*_, for instance, the Total Sum Scaling (TSS) and median-of-ratio (RLE)[9]. A large number of studies have demonstrated that different normalization methods will have a significant impact on the DE analysis, thus the choice of estimate method needs careful consideration. We chose to use the cumulative-sum-scaling (CSS)[11] in our model since it effectively controls the FDR. The methodology can be formulated as below (use RNA as an example):

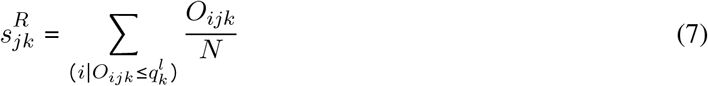

where 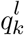 is the *l*th quantile of sample *k* and *N* is an appropriately chosen normalization constant.

### 3.4 Conditional estimation for dispersion parameters

We use the basic idea in DESeq2[9] to estimate the dispersion parameter *ϕ*_*ijk*_. Specifically, we first get a gene-wise estimation of the dispersion parameter, then shrink it towards the common trend. This approach leverages the information between genes effectively and stabilizes the estimation.

We get the estimation of gene-wise dispersion parameter by maximizing the Cox-Reid adjusted likelihood:

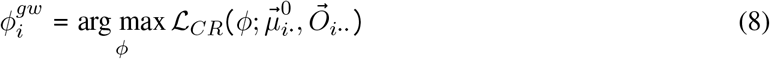

where 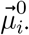 is an initial fit of the mean parameter using generalized negative binomial regression.

However, there is something different in our conditional negative binomial model. Since the observed gene count *C*_*ijk*_ can be also view as a random variable, the marginal distribution of *O*_*ijk*_ is hard to acquire. Thus, instead of maximizing the marginal distribution of *O*_*ijk*_, we maximizing the conditional distribution:

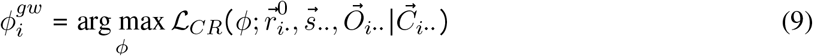

To achieve this, we need to first get the “normalized data” by dividing the MTX data with the paired MGX data (Our approach to handling zeros can be seen in the next section), then get an initial estimation of the mean parameter of normalized data using the negative binomial regression. This process can also be viewed as getting a crust estimation of the expression rate 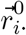. Next, we plug in the estimated normalized factors and expression rate parameter and maximize the likelihood function to get the estimate of the dispersion parameter.

A similar mean-dispersion trend is observed in MTX and MGX data. To use information between genes, we adopted the methods in DESeq2[9] and in Voom[10]. We first learn a parametric curve by regressing the gene-wise dispersion estimates onto the mean, then form a posterior for the dispersion from the likelihood function and the logarithmic prior. The value that maximizes the posterior is the final estimate of dispersion (MAP).

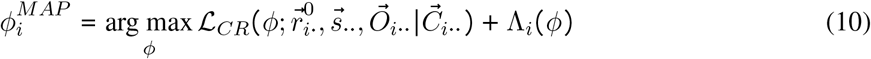

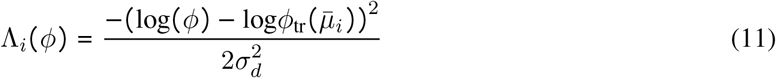

where 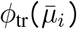 is the predicted dispersion based on the parametric curve, and 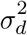 is the estimated prior variance.

### 3.5 Pre-filtering of technical zeros

A critical challenge in analyzing MGX and MTX data is the excess of zeros in read counts. We need to handle these zeros appropriately in order to give a robust and accurate inference. An ideal approach should filter the technical zeros while retaining biological zeros.

Previous work[14] has proposed some methods for pre-filtering technical zeros: ‘lenient’ filtering, ‘semi-strict’, and ‘strict’ filtering. They suggest using ‘strict’ filtering, which filters samples if either their MTX or gene-copy value was zero. However, the ‘strict’ filtering eliminates all zeros in data and fails to use the information in biological zeros. Here, we propose a different approach to pre-filter zeros in paired MGX-MTX data.

Our approach is to filter all samples with DNA=0 (no matter RNA=0 or not). The intuition behind this is simple:

‐ If the corresponding RNA!=0, we believe the zero in DNA is due to errors in experiments since the gene shouldn’t be lost if its transcript exists.
‐ If the corresponding RNA=0, this sample contains no information for expression rate and we shouldn’t use it.

However, if a sample has RNA=0 while DNA!=0, we tend to believe that the gene are biologically not expressed, and we keep these samples for further analysis.

### 3.6 Estimation of fold changes parameters

We get an initial estimate of Fold Changes (FCs) using a similar approach in DESeq2[9], by postulating a zero-centered normal prior for the FCs and maximizing the sum of the logarithmic likelihood of the GLM and the logarithm of the prior density. We called this result Bayesian shrunk FCs:

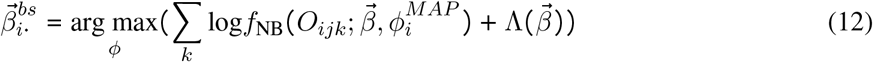

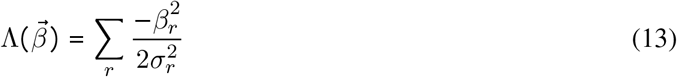

where 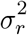 is the estimated prior width.

### 3.7 Likelihood ratio test for statistical inference

We use the likelihood ratio test to do statistical inference on the fold change parameter:

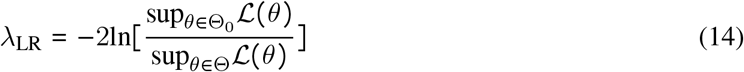

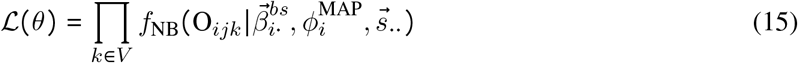

Where *V* represents the set of samples that are not filtered, *f*_*NB*_ refers to the mass function of the negative binomial distribution. According to Wilk’s theorem[20], the test statistics converge asymptotically to being *χ*^2^-distributed:

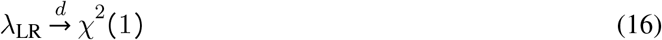

Since only one degree of freedom is lost (the group information). The obtained p-values are adjusted using Benjamini-Hochberg (BH) Procedure to control the FDR.

### 3.8 Bootstrap for fold changes shrinkage

One disadvantage of DESeq2[9] is that it can’t control the FDR well. The over-dispersion nature of the microbiome makes it extremely noisy and sensitive to extreme values. The extremely large read counts may lead to an inaccurate estimation of the FCs, especially for non-DE genes. Though DESeq2[9] has taken an empirical Bayesian shrinkage method for FCs estimation (Bayesian shrunk FCs), it performs badly for genes with relatively small but significant FCs. Through some investigations of the real data, we found that non-DE genes which are incorrectly inferred as significant often have larger standard deviations of FCs parameters. Thus, we propose to use bootstrap to shrink the estimation of FCs.

Specifically, after obtaining an initial set of significant genes, we re-estimate the FCs of these genes and do inference again. We first use bootstrap to get 50 estimates of the Bayesian shrunk FCs and compute its standard deviation (sd). Then we shrink the FCs toward zero, it can be formulated as:

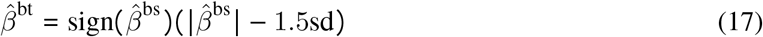

Using the bootstrap-shrunk FCs, we re-run the likelihood ratio test and obtain the final list of differentially expressed genes. It’s worth noting that:

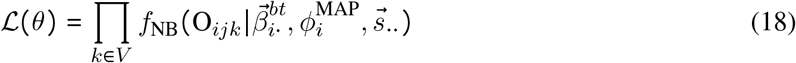

The likelihood is obtained using the new estimated FCs. It should be noted that the new significant genes should only be selected from the original set of significant genes. The bootstrap is only an approach to make our method more conservative. If the signal is weak in a dataset, we may choose not to run the bootstrap procedure to find more genes.

### 3.9 Simulation framework

The conditional negative binomial model can be formulated as below:

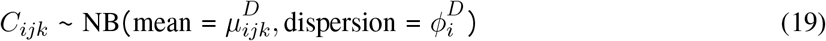

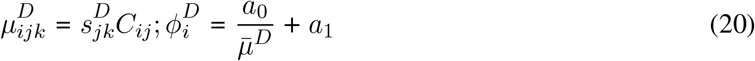

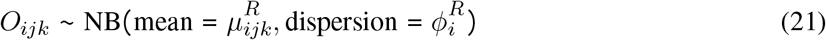

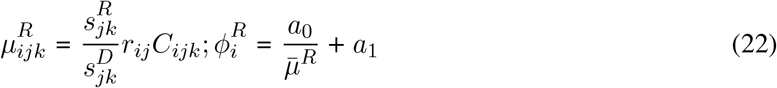

The model can simulate the relationship between MTX and MGX data. Thus, it is useful for bench-marking DE analysis that focuses on paired MGX-MTX data.

### 3.10 Performance evaluation

We evaluate the false discovery rate (FDR) and true positive rate (TPR) of our method using simulated datasets. A gene is defined to be biologically differentially expressed if there is a difference in the expression rate parameter *r*_*ij*_ between different groups. The change in RNA abundance introduced by underlying DNA is not considered a DE signal. A gene is recognized as significant if its p-value is smaller than 0.05 after the Benjamini-Hochberg correction[21]. The FDR and TPR can be computed using the following formula:

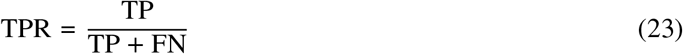

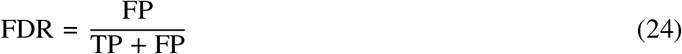

## 4 Results

### 4.1 Characteristics of metatranscriptomics data with sample-paired metagenomics data

The paired MGX-MTX data have some unique characteristics. Regarding the zeros in the paired data, there are four combinations in total: DNA!=0 and RNA!=0, DNA=0 and RNA!=0, DNA!=0 and RNA=0, DNA=0 and RNA=0. We plotted a heatmap to show the distribution of these four kinds of combinations using 500 genes (with zero proportion < 90%) in the ZOE dataset. From (a) in Figure 1, a high proportion of entries had both DNA=0 and RNA=0, while entries that had only one kind of data (MGX or MTX) equaled zero also took up a certain proportion. The lower the mean RNA abundance was, the higher proportion of zeros would occur in one gene. The distributions of these zeros had little correlation with the group information (caries-free).

**Figure 1:**
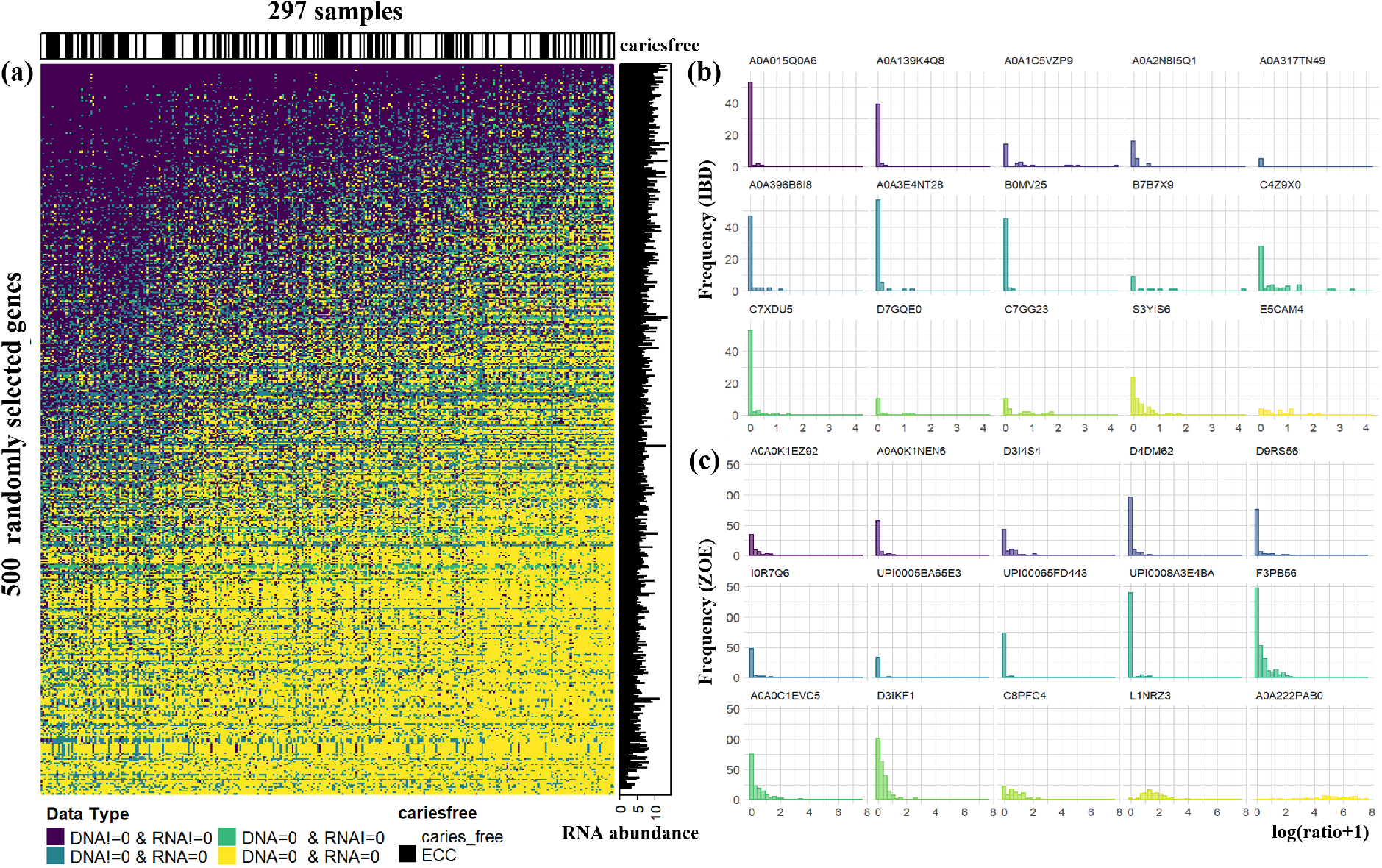
The distribution of zero types and RNA/DNA ratios in paired metatranscriptomics and metagenomics data. The ratios are plotted after filtering samples with DNA=0. (a) The distribution of different paired zero types in 500 randomly selected genes in the ZOE dataset. (b) The histogram of the RNA/DNA ratios in 15 randomly selected genes in the IBD dataset. (c) The histogram of the RNA/DNA ratios in 15 randomly selected genes in the ZOE dataset.

We also plotted the RNA/DNA ratios using only samples with DNA!=0. We selected 15 genes from IBD and ZOE 2.0 data sets for illustration. According to (b), (c) in Figure 1, there was not a fixed distribution of the RNA/DNA ratios, which complicated the direct analysis of ratios. Most ratios were less than 10 (log(ratio+1)<2), which provided some guidance for selecting expression rate parameters in the simulation.

### 4.2 Conditional estimation effectively eliminates the variation caused by underlying variation of gene copy number

When estimating the dispersions of the conditional negative binomial model, the underlying DNA information is used. The DNA information explains some of the variations of the RNA abundance, thus resulting in a reduced estimation of dispersions, which can potentially improve the power of differential expression analysis.

We plotted the dispersions versus mean RNA abundance using random 750 genes from the IBD dataset. The nonconditional dispersions were acquired only using the RNA information. Specifically, we first got a crust estimation of the mean parameter of RNA by fitting a negative binomial regression model. Next, we plugged in the estimated mean and estimated dispersions by maximizing the likelihood function. The conditional dispersions were acquired using a different approach. We first got an RNA/DNA ratio matrix (didn’t use the entries with DNA=0) and then estimated the mean of each group using a linear model. The mean could be viewed as a coarse estimation of the expression rates in different groups. Then we computed the mean of RNA abundance by multiplying the expression rate with DNA abundance and size factors. The dispersions were obtained again by maximum likelihood estimation.

From Figure 2, we observed a significant drop between the nonconditional dispersions and conditional dispersions. There was still a weak mean-dispersions trend according to the zoomed plot on the left side. The reduced dispersions also demonstrated the correctness of our model. Since if there wasn’t such a dependency between DNA and RNA, we were less likely to get a reduced estimation of dispersions.

**Figure 2:**
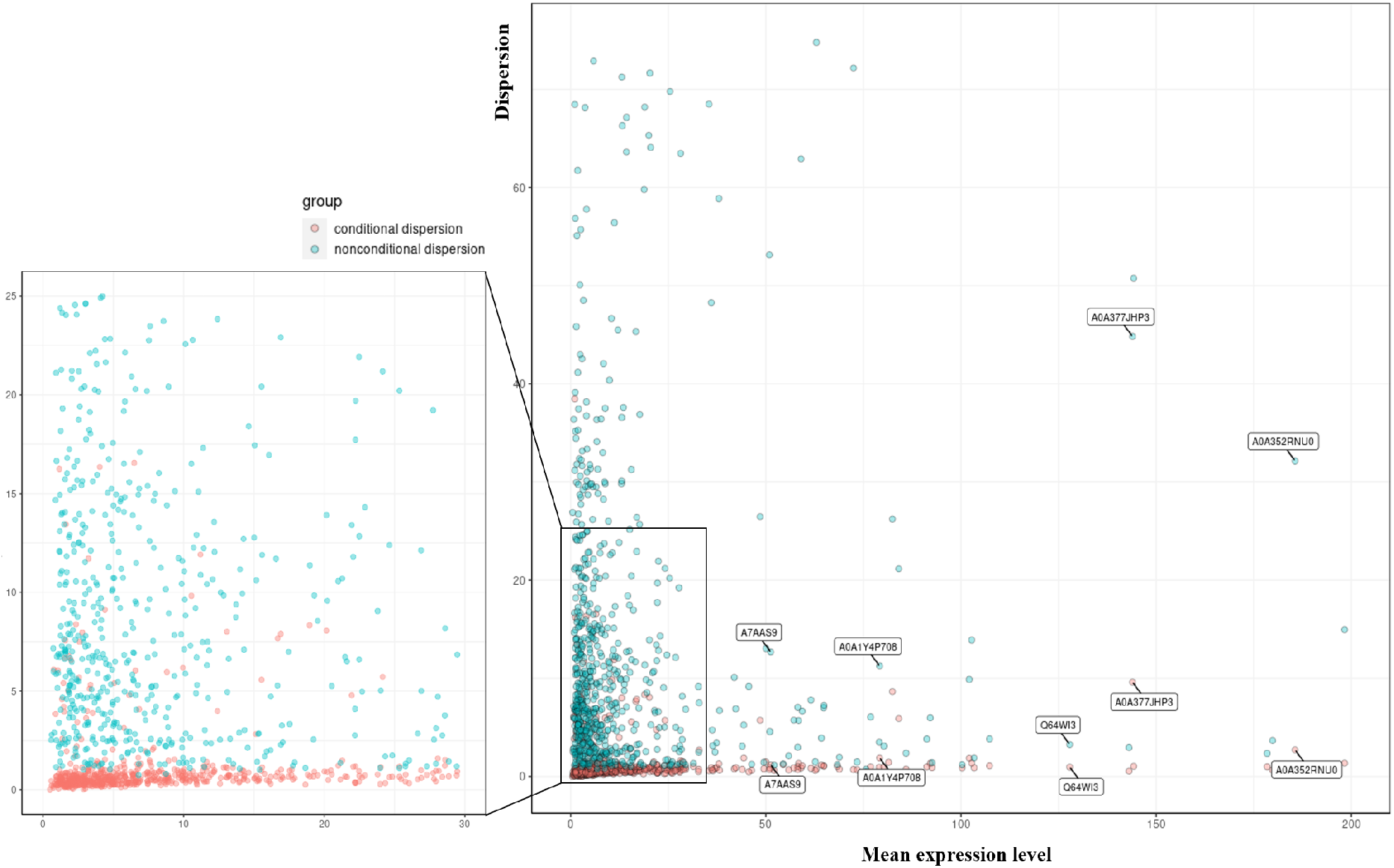
Scatter plot of the dispersions of 750 randomly selected genes in the IBD dataset. The conditional dispersions have the same mean expression level as their corresponding nonconditional dispersions. Some genes are labeled to show the relationship between their conditional and nonconditional dispersions.

### 4.3 Pre-filtering of technical zeros can improve model sensitivity and precision

We illustrated three different approaches to pre-filter zeros in paired MGX-MTX data in Figure 3. Our approach corresponded to moderate filtering. The strict filtering eliminated all zeros in the data and thus failed to use the information in biological zeros. The loose filtering treated the expression rates of entries with DNA=0 and RNA=0 as zeros, which may misuse the information in the data.

**Figure 3:**
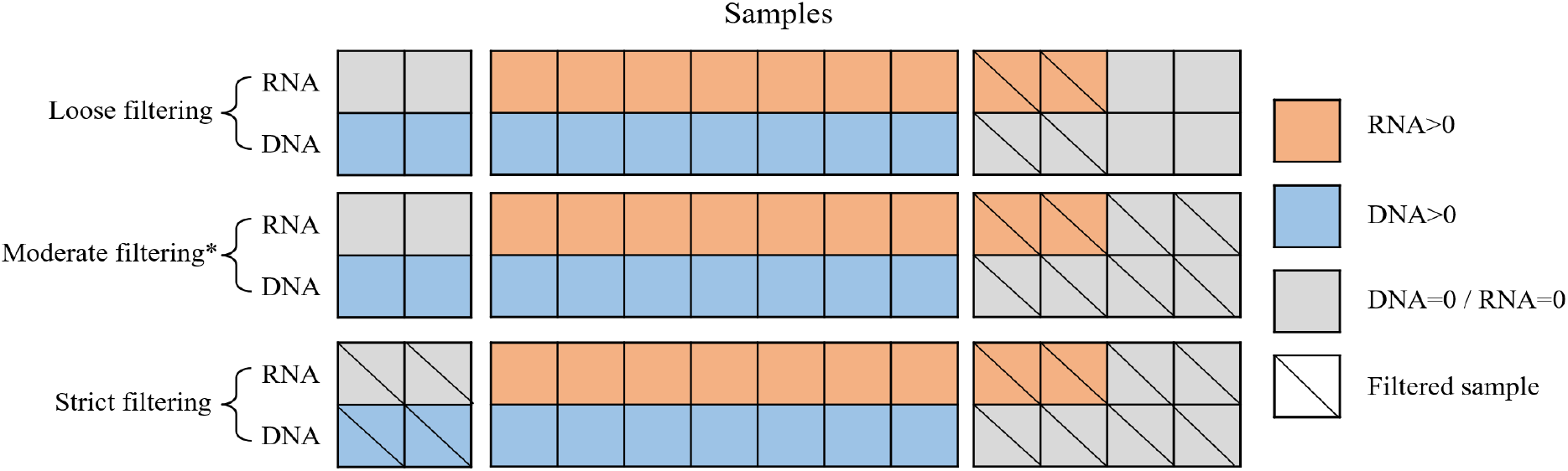
Graphical illustration of three different approaches to pre-filtering zeros.

We ran two simulations to demonstrate the effectiveness of our moderate filtering. The definition of simulations with/without confounding can be found in the Simulation results. We compared three different filtering methods across different sample sizes, with a fold change of 3. We could see that the loose filtering faced the problem of being over-conservative, while the strict filtering would inflate FDR in some cases. In comparison, Our moderate filtering could keep a high power while controlling the FDR well.

### 4.4 Simulation results

We simulated datasets of 300 genes with 10% proportion differential expressed genes using the framework mentioned in the Methods part. When selecting the mean, dispersions, and expression rate, we kept them at the same level as the parameters of the real data.

We simulated datasets in two different settings. The first simulation was generated when there was confounding. According to our model, the mean parameter of RNA count was affected by two parameters: the expression rate *r*_*ij*_ and the DNA abundance *C*_*ijk*_. In this case, we multiplied the expression rate of 10% genes by a certain fold change. These genes represented genes that were biologically differentially expressed. At the same time, we multiplied the paired DNA count of another 10% genes by the same fold change, and these genes were not considered as differentially expressed. The second simulation was generated when there was no confounding, which meant all changes in RNA abundance were due to changes in their expression rates.

Under each circumstance, we compared the performance of different methods under varying sample sizes (25-25, 50-50, 100-100), and different fold changes (2,3,4) were used to generate negative binomial count across two samples. The up-regulated group was chosen randomly. We repeated each simulation 30 times.

We compared four methods when there was confounding. The MTX here represented the method suggested in [14], we used its default parameters when running the simulation. The LN-dna method used a linear model to analyze the logarithmically transformed data. It leveraged DNA information by adding DNA as a covariate in the linear model. We also reported the performance of DESeq2 in the simulation. Please note that it was unfair to compare our method with traditional methods like DESeq2 when there was confounding since it was impossible for them to distinguish confounding without using the DNA information. We reported its performance just for showing the advantages of methods that used DNA information since they could identify genes that were biologically differentially expressed. The incapacity of traditional methods to identify changes introduced by underlying DNA abundance was easy to see according to Figure 5. For the comparison of three other methods, our method had the consistently highest power across all different the sample sizes and fold changes. Meanwhile, except for slight inflation of FDR when sample size was 25-25 and fold change is 2 (the weakest signal), our method controlled the FDR very well.

**Figure 4:**
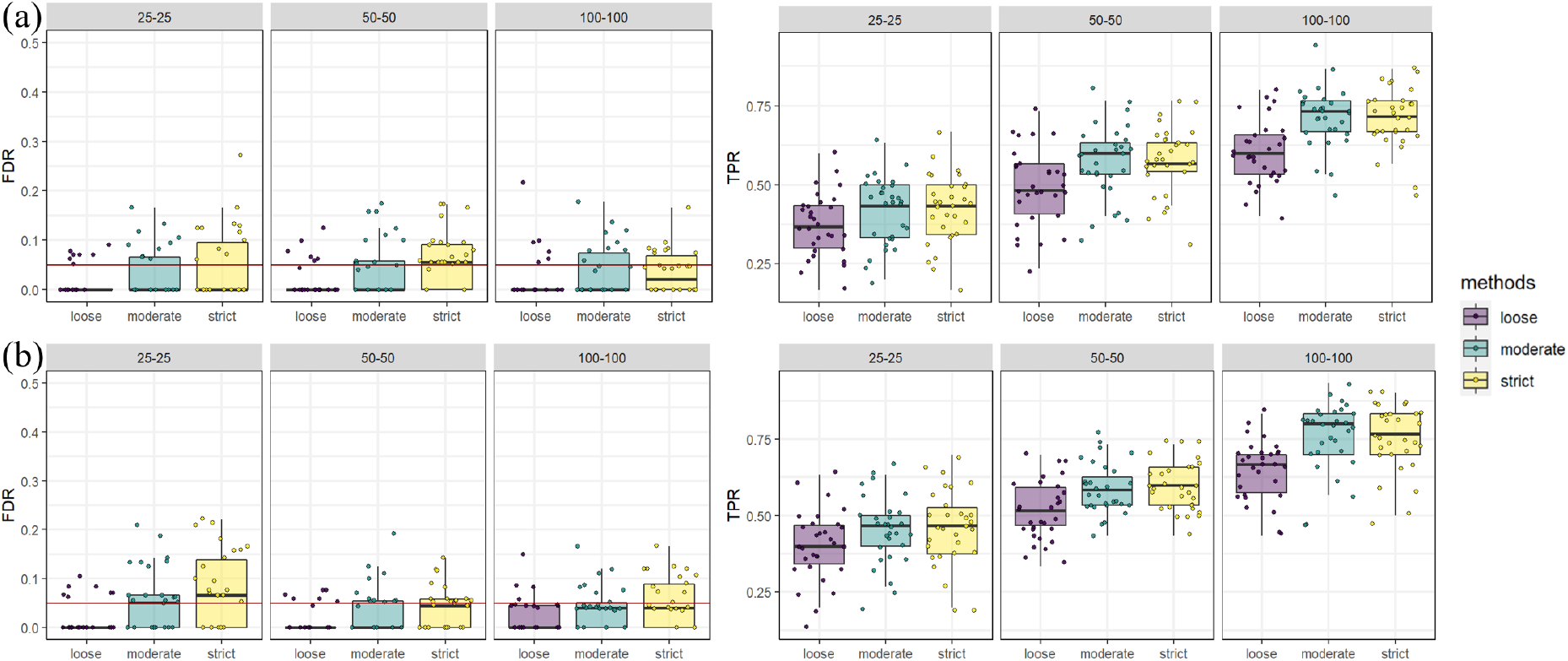
Simulation results comparing different zero pre-filtering methods. (a) There is confounding (b) There is no confounding

**Figure 5:**
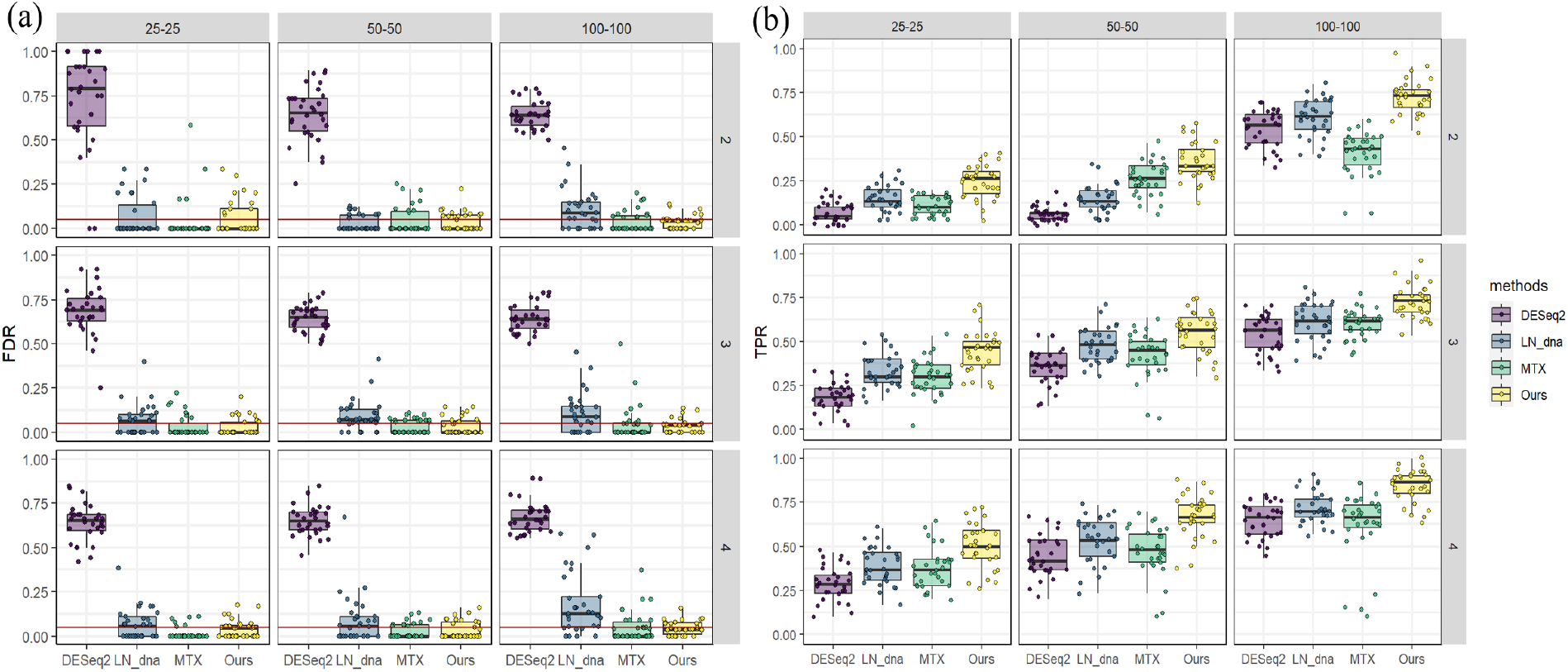
Simulation results when there is confounding. There are a total of 9 different combinations of sample size and fold change. For example, “50-50” means 50 samples in each group; “3” means a fold change of 3. (a) False discovery rate of different methods; the red line indicates a significance level of 0.05 (b) True positive rate of different methods.

When there was no confounding, we compared our model with a total of 7 other methods, including six traditional ones and the MTX method. Again, our methodology showed the highest power across different sample sizes and fold changes while controlling the FDR well (except in the case with the weakest signal). Methods that leveraged DNA information had an overall better performance than the traditional methods, which demonstrated the necessity of using DNA information.

### 4.5 Real data based simulation for controlling Type I Error

We demonstrated the performance of controlling Type I Error in our method using the IBD and ZOE 2.0 datasets. Although the real data are helpful to verify how well the theory fits reality, it faces the problem of not knowing the underlying truth, thus is difficult to evaluate the performance using traditional methods. To overcome this problem, we first carried out a negative control experiment to see how well our methodology controlled the FDR in real datasets. We randomly selected 2000 genes (with zero proportion < 90%) and randomly assigned the samples into two groups. Due to the randomness of the assignment, no differentially expressed genes were expected. Under these circumstances, all the significant genes found by DE methods were false positives. We repeated the simulation 30 times using each of the datasets. According to Figure 7, in the IBD dataset, the DESeq2 found the most false positives, which indicated that DESeq2 didn’t controll FDR well in the IBD dataset. The edgeR also consistently identified more significant genes compared with other methods. Although our model also found more significant genes than some methods (ALDEx2, metagenomeSeq, etc.), it kept the false positive calls ≤ 2 in all 30 simulations (with a total of 2000 genes). In the ZOE 2.0 datasets, although edgeR and DESeq2 still found slightly more significant genes, all other methods controlled the false positive very well. The negative control experiment demonstrated the ability of our methodology to control the false positives.

**Figure 6:**
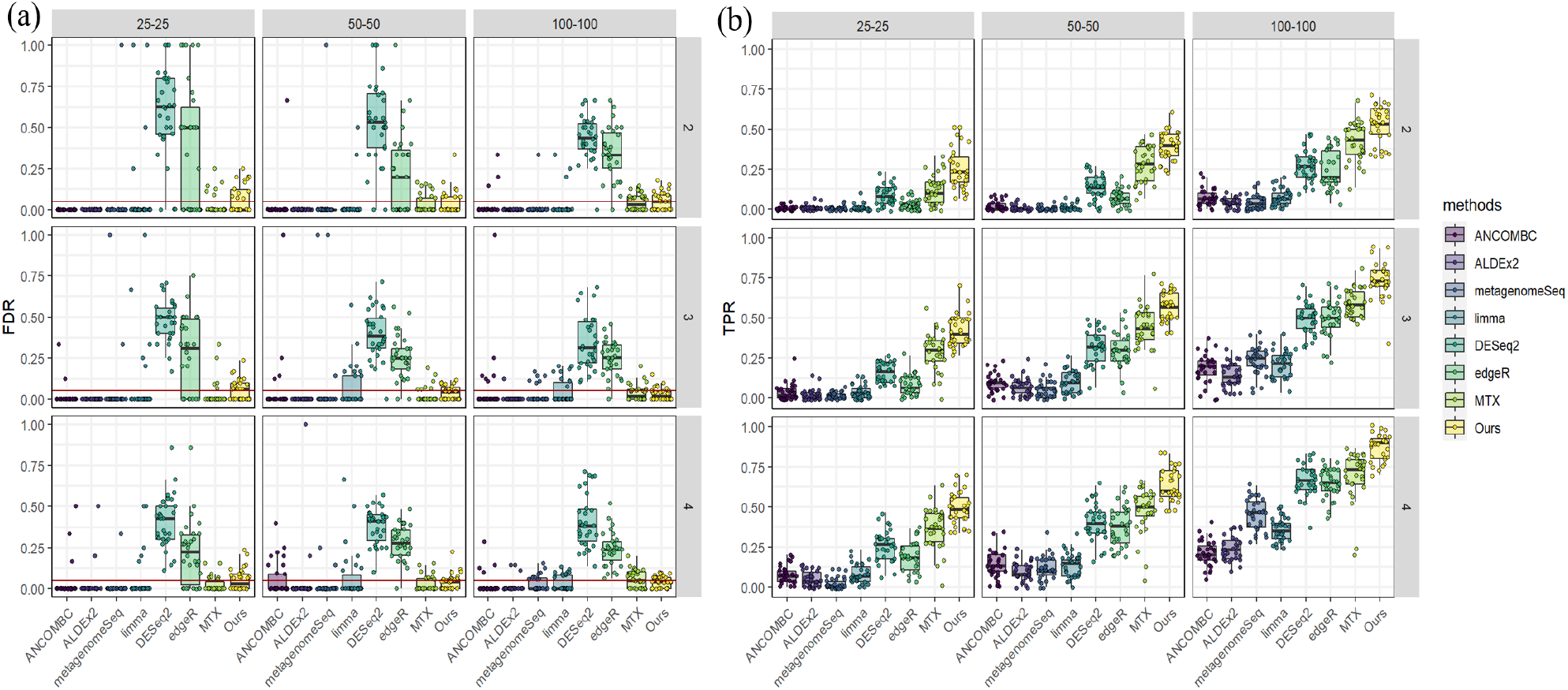
Simulation results when there is no confounding. (a) The false discovery rate of different methods; the red line indicates a significance level of 0.05 (b) True positive rate of different methods.

**Figure 7:**
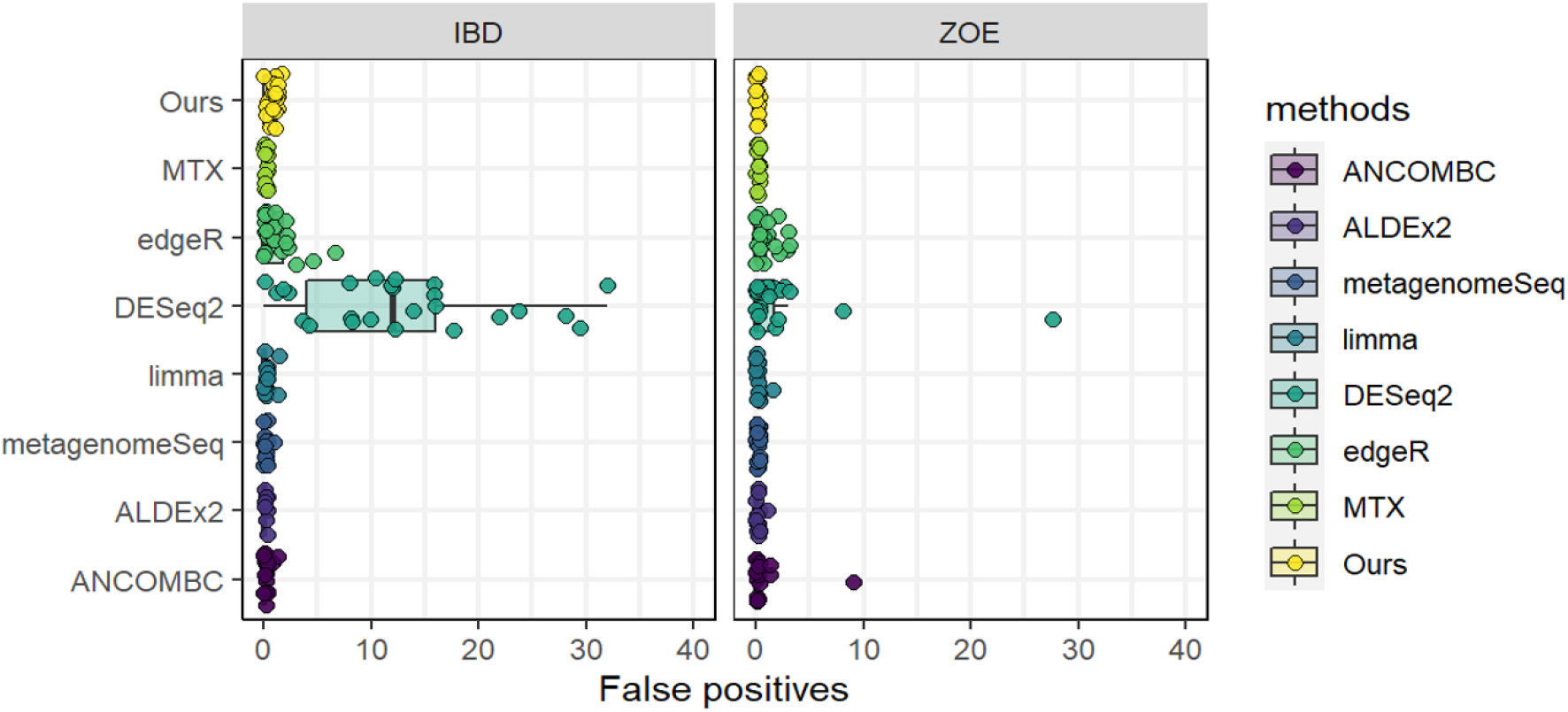
False positives of different methods in negative control experiment.

### 4.6 Application in oral microbiome data

Using the method we developed for DE analysis, We reported several significant genes identified in the ZOE 2.0 datasets of which the expression activity was associated with the early childhood caries (ECC) phenotype. The ZOE 2.0 datasets have 210158 genes and 297 samples, with a 49% (147/297) ECC prevalence. To discover more genes, we didn’t run the bootstrap procedure and finally found 14 genes that were differentially expressed between different ECC phenotypes. The Uniref90 ID, corresponding proteins, and microorganisms were summarized in Table 2. Among all these microorganisms, Corynebacterium matruchotii[22], Capnocytophaga[23], Prevotella nigrescens[24], Tannerella[25], Veillonella parvula[26] and Fusobacterium nucleatum[27] are known to be related to dental health. Some articles also support Streptococcus oralis[28], Streptococcus anginosus[29], and Streptococcus cristatus[30] are related to caries. The result further demonstrated the feasibility of our method.

**Table 1:** Summary of notations

| Notations | Description |
| --- | --- |
| $i$ | gene index, $i = 1, 2, 3, \dots, n$ |
| $j$ | group index, $j = 1, 2, 3, \dots, g$ |
| $k$ | sample index, $k = 1, 2, 3, \dots, m_j$ |
| $C_{ij}$ | expected abundance of gene $i$ in group $j$ |
| $O_{ij}$ | expected abundance of RNA $i$ in group $j$ |
| $s_{jk}^D$ | library size of sample $k$ for gene in group $j$ |
| $s_{jk}^R$ | library size of sample $k$ for RNA in group $j$ |
| $C_{ijk}$ | Observed abundance of gene $i$ of sample $k$ in group $j$ |
| $O_{ijk}$ | Observed abundance of RNA $i$ of sample $k$ in group $j$ |

**Table 2:** Summary of significant genes

| UnirefID | Protein | Organism |
| --- | --- | --- |
| D4FR29 | 50S ribosomal protein L13 | Streptococcus oralis |
| E0DDK1 | Holin | Corynebacterium matruchotii |
| L1P0V7 | Response regulator receiver domain protein | Capnocytophaga sp. oral taxon 332 |
| L1PWD4 | DUF6268 domain-containing protein | Capnocytophaga sp. oral taxon 326 |
| U2Z4S8 | Uncharacterized protein | Streptococcus anginosus group |
| U2ZPB1 | Uncharacterized protein | Streptococcus anginosus T5 |
| V8CH43 | Uncharacterized protein | Prevotella nigrescens CC14M |
| W2CNH7 | Antitoxin Phd_YefM | Tannerella sp. oral taxon BU063 |
| A0A0C1H2V0 | Transposase | Morococcus cerebrosus |
| A0A0F2CG03 | Peptide deformylase | Streptococcus cristatus |
| A0A134C019 | stress response membrane protein | Veillonella parvula |
| A0A154TIE5 | 50S ribosomal protein L35 | Neisseria flavescens |
| A0A1B2Z6U0 | Protein kinase domain-containing protein | environmental samples |
| A0A2C6C3R2 | MarR family transcriptional regulator | Fusobacterium nucleatum |

## 5 Discussion

Metatranscriptomics is essential for revealing the functional activities and gene expression patterns of a complex assemblage of microbes. However, due to the dynamic nature of microbial communities, the transcripts abundance is affected by many other factors like taxon abundance, which complicates downstream analysis. Differential expression analysis is an important task when analyzing metatranscriptomics data, which can identify genes that are related to disease status. The taxon-specific scaling may lose power when some genes can’t be assigned to a specific taxon. Therefore, when the paired MGX-MTX data is available, consideration of DNA abundance in the DE analysis allows the identification of DE genes with differential transcriptional activities that are not due to the taxa abundance change.

Here, we propose a differential expression analysis method for metatranscriptomics with paired metagenomics data. The over-dispersion, zero-inflation nature of metatranscriptomics data, as well as the dependency between RNA and underlying DNA, have posed challenges to the DE analysis. Our conditional negative binomial model leverages DNA information to improve the power of DE analysis and can yield a biologically meaningful result. By considering the underlying DNA abundance, our method can successfully identify the changes in RNA abundance which are due to gene copy number variation, thus circumventing spurious community DE signal. The over-dispersion characteristic of MTX complicates corresponding statistical modeling and may reduce the power of DE analysis. We discover that by relating DNA abundance with the mean of RNA abundance distribution, we get a reduced estimation of dispersions which can improve the power of DE analysis. The discovery also demonstrates the appropriateness of the conditional model, by indicating a strong relationship between DNA and RNA abundance. Zero inflation is also a troublesome characteristic which is more complicated in paired MTX and MGX data. To address this problem, we suggest using “moderate filtering” to deal with the zeros in paired data. This filtering method is a tradeoff between “strict filtering” and “loose filtering”, and achieved the best performance in simulations.

Particularly in simulation, we proposes a conditional negative binomial framework for generating paired MGX-MTX data, which is helpful for the evaluation of differential expression methods that directly model the paired data. This framework considers the dependency between RNA abundance and DNA abundance, which is difficult to take into account in traditional simulation methods. The simulation studies have demonstrated the superiority of our DE analysis method, which can maintain a high power controlling the FDR well. In the application to the ZOE 2.0 dataset, we find a total of 14 differential expressed genes, including genes belonging to the microbial species that are previously known associated with oral diseases, which further showed the feasibility of our method.

Despite the advantages, our work also has some limitations. Firstly, our method doesn’t consider the composition nature of the microbiome, which will also affect the DA/DE analysis of microbiome communities. Meanwhile, we take a relatively simple approach to deal with the zeros in the data, which may lose information and reduce power. Techniques like imputation or more complicated modeling of zeros are needed to address this problem. Also, a new synthetic data generation framework that considers the effects of other factors like species prevalence is needed for a more comprehensive evaluation.

In conclusion, our differential expression analysis method offers researchers a new tool that directly considers the underlying metagenomics data. By leveraging DNA information, our method can yield more biologically meaningful results, thus providing new chances for in-depth investigation of the functional abilities of the dynamic microbial communities.

## References

[1] Katherine R Amato. “An introduction to microbiome analysis for human biology applications”. In: American Journal of Human Biology 29.1 (2017), e22931.

[2] Christopher Quince et al. “Shotgun metagenomics, from sampling to analysis”. In: Nature biotechnology 35.9 (2017), pp. 833–844.

[3] Marco JL Coolen and William D Orsi. “The transcriptional response of microbial communities in thawing Alaskan permafrost soils”. In: Frontiers in microbiology 6 (2015), p. 197.

[4] Eric A Franzosa et al. “Relating the metatranscriptome and metagenome of the human gut”. In: Proceedings of the National Academy of Sciences 111.22 (2014), E2329–E2338.

[5] Mary Ann Moran. “Metatranscriptomics: eavesdropping on complex microbial communities”. In: Microbe 4.7 (2009), p. 7.

[6] Stavros Bashiardes, Gili Zilberman-Schapira, and Eran Elinav. “Use of metatranscriptomics in microbiome research”. In: Bioinformatics and biology insights 10 (2016), BBI–S34610.

[7] Abhishek Kaul et al. “Analysis of microbiome data in the presence of excess zeros”. In: Frontiers in microbiology 8 (2017), p. 2114.

[8] Mark D Robinson, Davis J McCarthy, and Gordon K Smyth. “edgeR: a Bioconductor package for differential expression analysis of digital gene expression data”. In: bioinformatics 26.1 (2010), pp. 139–140.

[9] Michael I Love, Wolfgang Huber, and Simon Anders. “Moderated estimation of fold change and dispersion for RNA-seq data with DESeq2”. In: Genome biology 15.12 (2014), pp. 1–21.

[10] Charity W Law et al. “voom: Precision weights unlock linear model analysis tools for RNA-seq read counts”. In: Genome biology 15.2 (2014), pp. 1–17.

[11] Joseph N Paulson et al. “Differential abundance analysis for microbial marker-gene surveys”. In: Nature methods 10.12 (2013), pp. 1200–1202.

[12] Greg Gloor. “ALDEx2: ANOVA-Like Differential Expression tool for compositional data”. In: ALDEX manual modular 20 (2015), pp. 1–11.

[13] Huang Lin and Shyamal Das Peddada. “Analysis of compositions of microbiomes with bias correction”. In: Nature communications 11.1 (2020), pp. 1–11.

[14] Yancong Zhang et al. “Statistical approaches for differential expression analysis in metatranscriptomics”. In: Bioinformatics 37.Supplement_1 (2021), pp. i34–i41.

[15] Heiner Klingenberg and Peter Meinicke. “How to normalize metatranscriptomic count data for differential expression analysis”. In: PeerJ 5 (2017), e3859.

[16] Guillem Salazar et al. “Gene expression changes and community turnover differentially shape the global ocean metatranscriptome”. In: Cell 179.5 (2019), pp. 1068–1083.

[17] Francesco Beghini et al. “Integrating taxonomic, functional, and strain-level profiling of diverse microbial communities with bioBakery 3”. In: Elife 10 (2021), e65088.

[18] Kimon Divaris et al. “Cohort profile: ZOE 2.0—a community-based genetic epidemiologic study of early childhood oral health”. In: International Journal of Environmental Research and Public Health 17.21 (2020), p. 8056.

[19] Jason Lloyd-Price et al. “Multi-omics of the gut microbial ecosystem in inflammatory bowel diseases”. In: Nature 569.7758 (2019), pp. 655–662.

[20] Samuel S Wilks. “The large-sample distribution of the likelihood ratio for testing composite hypotheses”. In: The annals of mathematical statistics 9.1 (1938), pp. 60–62.

[21] Yoav Benjamini and Yosef Hochberg. “Controlling the false discovery rate: a practical and powerful approach to multiple testing”. In: Journal of the Royal statistical society: series B (Methodological) 57.1 (1995), pp. 289–300.

[22] Bruce J Paster et al. “Bacterial diversity in human subgingival plaque”. In: Journal of bacteriology 183.12 (2001), pp. 3770–3783.

[23] Anne Jolivet-Gougeon et al. “Antimicrobial treatment of Capnocytophaga infections”. In: International journal of antimicrobial agents 29.4 (2007), pp. 367–373.

[24] Catalina-Suzana Stingu et al. “Association of periodontitis with increased colonization by P revotella nigrescens”. In: Journal of investigative and clinical dentistry 4.1 (2013), pp. 20–25.

[25] Sasanka S Chukkapalli et al. “Chronic oral infection with major periodontal bacteria Tannerella forsythia modulates systemic atherosclerosis risk factors and inflammatory markers”. In: Pathogens and disease 73.3 (2015), ftv009.

[26] GPA Bongaerts et al. “Was isolation of Veillonella from spinal osteomyelitis possible due to poor tissue perfusion?” In: Medical hypotheses 63.4 (2004), pp. 659–661.

[27] Paul E Kolenbrander et al. “Bacterial interactions and successions during plaque development”. In: Periodontology 2000 42.1 (2006), pp. 47–79.

[28] PM Corby et al. “Microbial risk indicators of early childhood caries”. In: Journal of clinical microbiology 43.11 (2005), pp. 5753–5759.

[29] David Beighton. “The complex oral microflora of high-risk individuals and groups and its role in the caries process”. In: Community dentistry and oral epidemiology 33.4 (2005), pp. 248–255.

[30] ACR Tanner et al. “Cultivable anaerobic microbiota of severe early childhood caries”. In: Journal of clinical microbiology 49.4 (2011), pp. 1464–1474.

